# Scarabaeidae larvae are neglected greenhouse gas sources in soils

**DOI:** 10.1101/713784

**Authors:** Carolyn-Monika Görres, Claudia Kammann

**Affiliations:** Department of Applied Ecology, Hochschule Geisenheim University, Geisenheim, Germany; Department of Soil Science and Plant Nutrition, Hochschule Geisenheim University, Geisenheim, Germany

## Abstract

A precise knowledge of the sink and source distributions of greenhouse gases (GHG) in regional and global carbon and nitrogen budgets, and of the processes governing them, is a necessary prerequisite for the development and assessment of climate change adaptation and mitigation strategies^1-3^. Certain soil-inhabiting Arthropoda groups are known producers of GHG, namely methane (CH_4_), but apart from termites, their emissions have never been studied in the field and quantified at different scales^4,5^. Here we report the first field GHG emission data of soil-dwelling Scarabaeidae larvae, focusing on pest insects in a temperate climate region (*Melolontha melolontha* and *M. hippocastani*). Variations in larval biomass explained variations in larval field CO_2_ and CH_4_ emissions well at the individual and site level. This correlation disappeared after transferring larvae from the field to a laboratory setting. We show that GHG emissions of soil-inhabiting Scarabaeidae larvae are comparable to those from termites, thus questioning the neglect of Scarabaeidae larvae in GHG flux research, and we demonstrate the importance of field-based emission estimates for soil biota.

## Introduction

A major challenge in climate change research is to fully understand and accurately quantify interannual to decadal variabilities in atmospheric GHG concentrations driven by natural processes^**1,6**^. Soils can act as both, important sinks and sources of atmospheric carbon dioxide (CO_2_), CH_4_, and nitrous oxide (N_2_O), but are also components in the global carbon and nitrogen budget with large uncertainty estimates^**1-3**^. In recent years, it has been proposed to reduce these uncertainties by shifting from an implicit to an explicit representation of soil biota in ecosystem and, ultimately, Earth system models^**7,8,9**^. However, a major constraint for developing such new biogeochemical models is the lack of field data, especially for soil biota groups other than microorganisms^**8,9**^. Soil biota can substantially influence the spatial and temporal variability of GHG sinks and sources in the field, but the majority of our current knowledge comes from laboratory experiments, is often controversial and has been limited to only a few regions and species. As a result, the magnitude of the effect of soil biota on the GHG sink and source capacity of soils remains poorly quantified. For example, termites are the only soil-inhabiting Arthropoda group, which has been explicitly considered as significant GHG source on a global and regional level thus far^**4,10**^. They are assumed to contribute about 1 – 3 % to the global annual CH_4_ budget, but the large variation in emission estimates in the literature (0.9 to 150 Tg CH_4_ y^-1^) underlines the uncertainty in the available data sets^**2,4,11**^.

We conducted the first GHG field measurement study on soil-inhabiting larvae of the Scarabaeidae family. Throughout the vegetation period of 2017, larvae of *Melolontha melolontha* (Common cockchafer) and *M. hippocastani* (Forest cockchafer) were excavated at randomly chosen plots (0.25 m^2^ each, excavation depth ∼50 cm) at six sites (Site 1 – Site 6) in west-central and southern Germany covering different larval developmental stages and activity levels, and vegetation types. Greenhouse gas emissions of individual larvae were measured directly in the field together with larval biomass and various environmental variables. One site was selected for a comparison of field- and laboratory-measured GHG emissions. For this, larvae were randomly separated into three groups. For one group, larval GHG emissions were measured directly in the field. The other two groups were transferred to the laboratory, where the larvae were kept individually in soil-filled containers with ample food supplies for 7 and 18 days, respectively, before GHG emission measurements were carried out (Methods). Larval biomass was the main variable used to describe inter- and intra-site GHG emission variability, and to subsequently model total larval plot-level CO_2_ and CH_4_ emissions using linear regression analysis on the pooled field dataset. Upscaled plot-level emission and biomass estimates represented the cumulated individual measurement values per plot standardized to 1 m^2^. To derive an annual European CO_2_ and CH_4_ emission estimate, the range of plot-level emissions was multiplied by the available literature value on area colonized by *Melolontha* spp. in Europe^**12**^. Individual larval N_2_O emission were not upscaled, due to their infrequent occurrence (Methods).

## Results

Gaseous carbon emissions of individual *Melolontha* spp. larvae showed a large inter- and intra-site variability which could not be explained by differences in soil temperature (range: 11.4 – 29.3 °C) and soil moisture (range: 3.2 – 32.7 vol%). There was a clear tendency for emissions to increase with larval biomass at the site level, especially for CO_2_ (Fig. 1). Average larval biomass ranged between 0.5 and 2.2 g larva^-1^. When pooling all data regardless of site and species, there was a strong positive correlation between CO_2_ and CH_4_ emissions (*r*_*s*_ = 0.76, p<0.001), and between larval biomass and CO_2_ emissions (*r*_*s*_ = 0.84, p<0.001). The correlation between larval biomass and CH_4_ emissions was less pronounced, but still significant (*r*_*s*_ = 0.68, p<0.001). The excavation depth of the larvae correlated negatively with larval biomass (*r*_*s*_ = −0.51, p<0.001), CO_2_ emissions (*r*_*s*_ = −0.58, p<0.001), and CH_4_ emissions (*r*_*s*_ = −0.48, p<0.001), respectively (Supplementary information S1). It could be seen as an indicator for larval access to fresh plant root material, which was higher the closer the larvae were to the soil surface, i.e. the lower the excavation depth was. Two-thirds of the larvae were found at 0 – 15 cm soil depth. Nitrous oxide emissions were only occasionally observed. Out of the 64 field larvae tested for N_2_O (sites 2 – 6), 13 individuals emitted significant amounts, ranging between 1.3 and 90.4 ng N_2_O h^-1^ larva^-1^.

**Figure 1:**
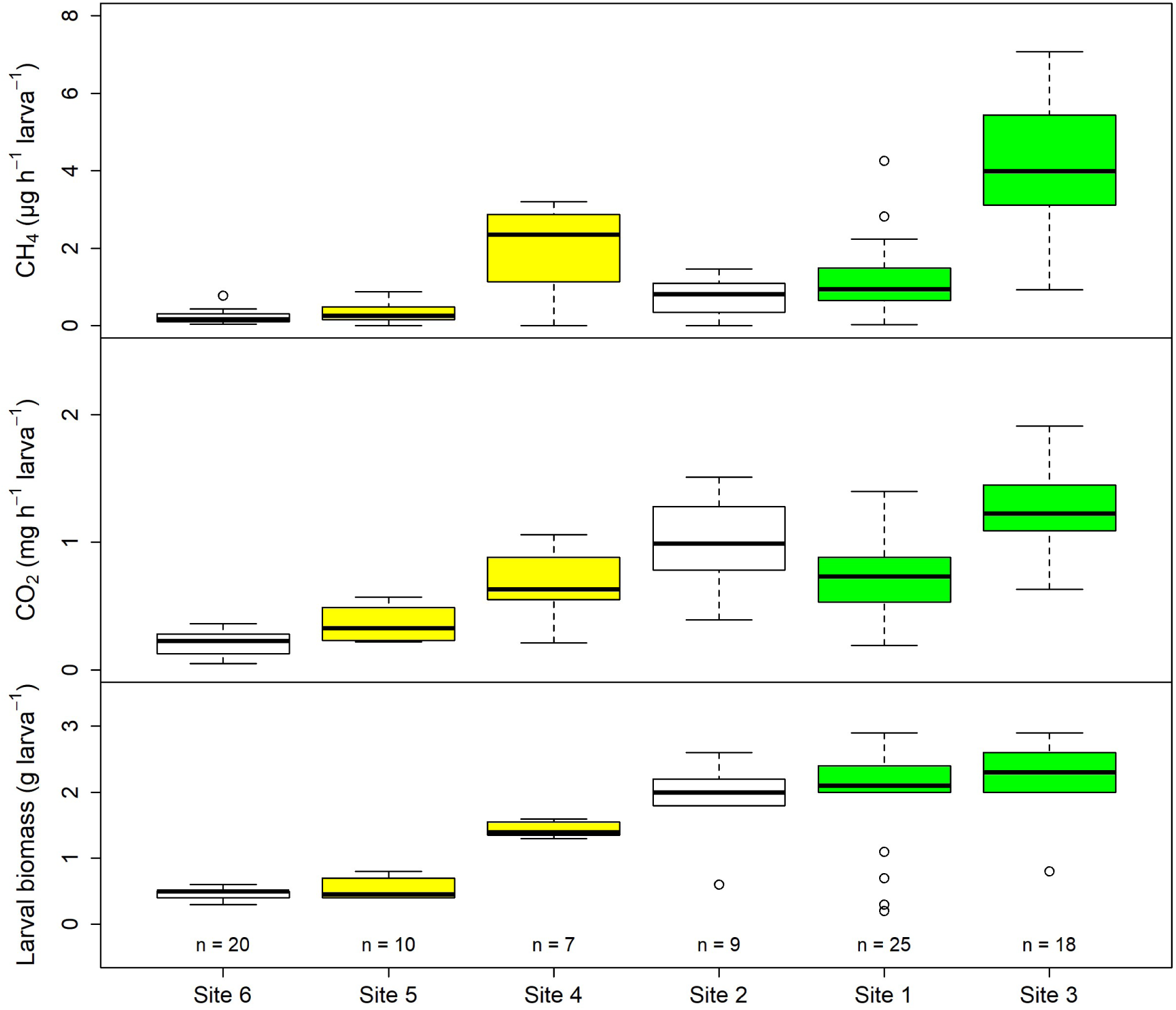
Direct CH_4_ and CO_2_ emissions, and larval biomass of individual *Melolontha* spp. larvae during field incubations. The box midline shows the median, with the upper and lower limits of the box being the 75th and 25th percentile, respectively. The whiskers extend up to 1.5 times from the box edges to the furthest data point within that distance. Sampling sites are sorted by average larval biomass in ascending order. The colours of the boxplots indicate the species (white = *Melolontha* spp., green = *M. melolontha*, yellow = *M. hippocastani*).

Since Scarabaeidae larvae need to reach a certain biomass to be able to pupate and since their food intake increases with size, larval biomass was a good proxy for larval age, sampling time, and food availability. Larval biomass could also be used to differentiate between species and to encode larval abundances at the plot scale. Larval abundances ranged between 4 and 68 larvae m^-2^ (Supplementary information S2). The correlation between gaseous carbon emissions and larval biomass persisted when upscaling emissions to the plot scale, and a large proportion of the inter- and intra-site emission variability could be explained by variations in total larval biomass (Fig. 2). Across all sites, CO_2_ emissions increased on average by 0.51±0.03 mg CO_2_ h^-1^ m^-2^ with each g total larval biomass increase (p<0.001). The relationship between CH_4_ emissions and total larval biomass was best fitted with a linear regression model including a polynomial term (Supplementary information S3). Plot-scale emissions ranged between 1.19 and 46.75 mg CO_2_ h^-1^ m^-2^, and 1.15 and 155.58 μg CH_4_ m^-2^ h^-1^ (excluding one plot at Site 4 with zero emissions) (Fig. 2). Based on these values and a literature value of 200,000 ha colonised by *Melolontha* spp. in Europe, gaseous carbon emissions from *Melolontha* spp. larvae alone were estimated to range between 10.42 and 409.53 kt CO_2_ yr^-1^, and 0.01 and 1.36 kt CH_4_ yr^-1^ in this region.

**Figure 2:**
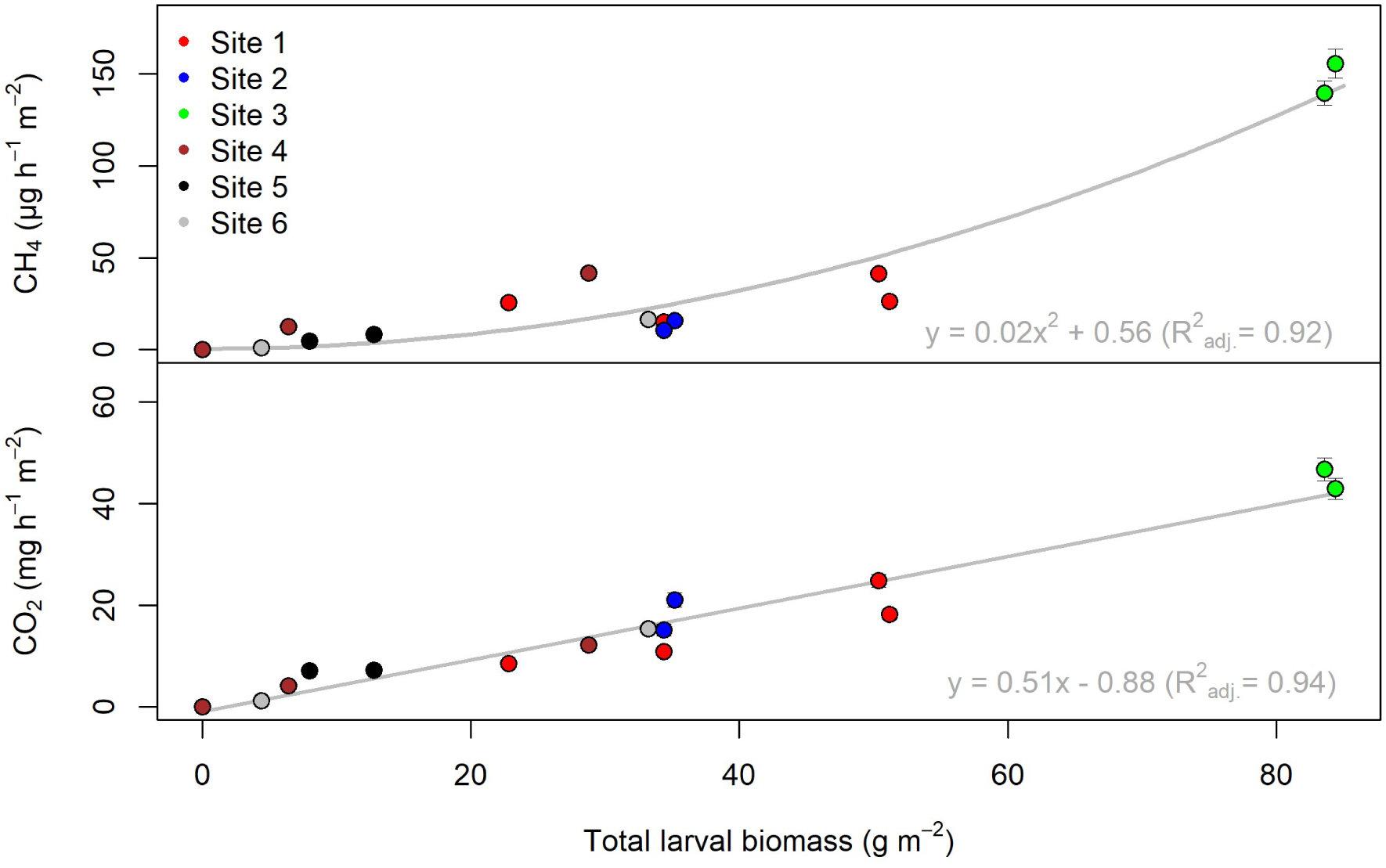
Cumulated larval CO_2_ and CH_4_ emission estimates from field measurements for each individual sampling plot in relation to total cumulated larval biomass grouped by sampling site. Error bars represent the propagated error of the individual larval emissons. Results of linear regression analysis using the complete dataset are given in the respective subfigure for CH_4_ and CO_2_.

A comparison of field and laboratory measurements from larvae excavated at Site 3 revealed a strong impact of laboratory conditions on CO_2_ and CH_4_ emissions. Despite no significant differences in larval biomass between the three measured groups (p=0.12) and ample food supply, overall emission strength and variability decreased rapidly with prolonged time at the laboratory (Fig. 3). In contrast to the field observations, no significant correlation between larval biomass and CO_2_ and CH_4_ emissions was found, respectively, after two and a half weeks in the laboratory. N_2_O emissions tended to be lower under laboratory conditions in comparison to field measurements as well; however, it was not possible to discern a statistically significant effect of the laboratory conditions on larval N_2_O emissions. Of the 65 larvae incubated in the laboratory, only 13 emitted N_2_O, with emissions ranging between 1.81 and 43.70 ng N_2_O h^-1^ larva^-1^ (Supplementary information S2).

**Figure 3:**
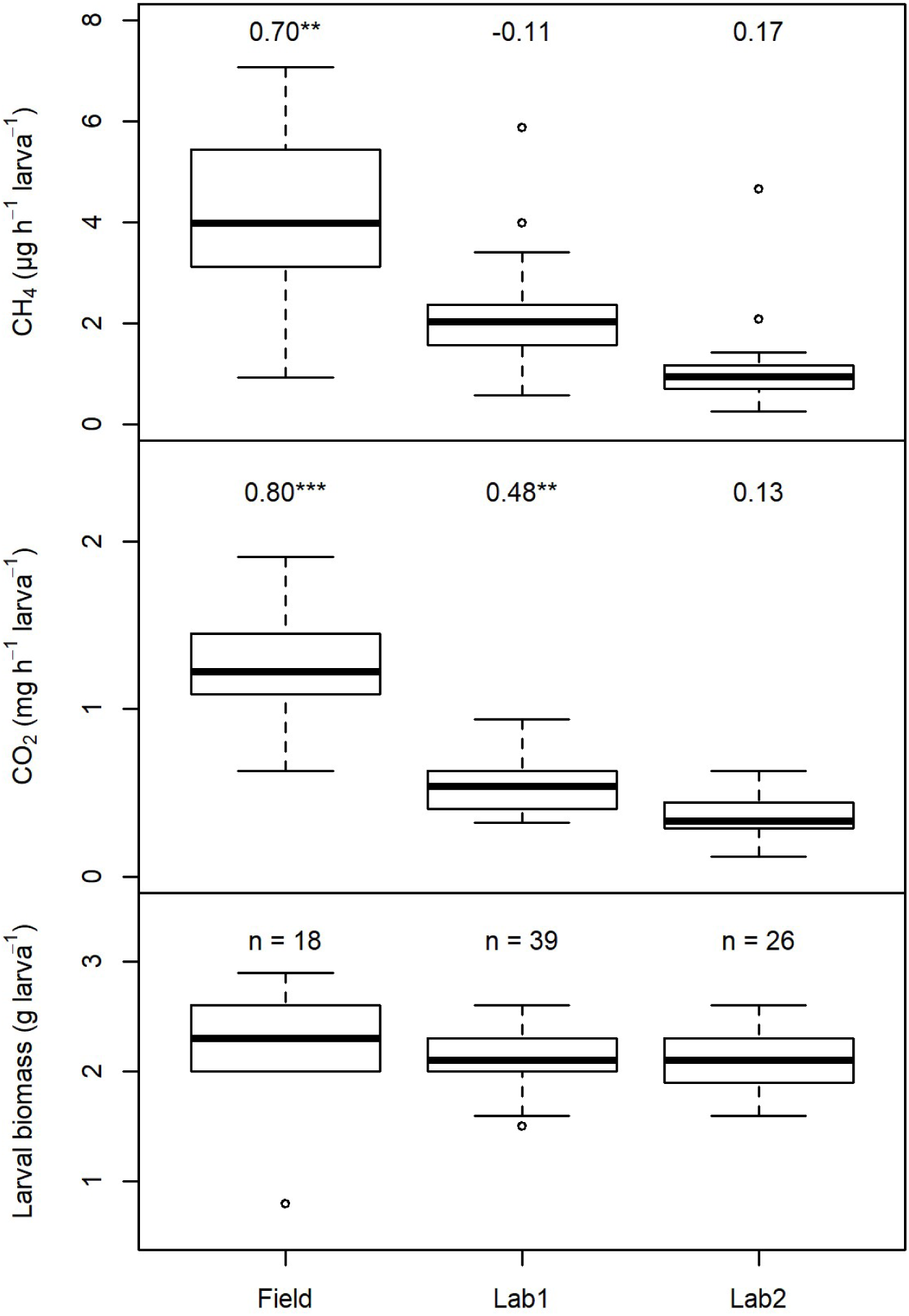
Comparison of direct *M. melolontha* larval CO_2_ and CH_4_ emissions from three different batches. All 83 larvae were excavated at sampling site 3 on 26.05.2017. Larval emissions of batch “Field” were sampled in the field directly after excavation. Batch “Lab1” and “Lab2” were kept 7 and 18 days in the laboratory, respectively, before emissions were measured. The boxplots for “Field” are identical to the boxplots for site 3 in Fig. 1. Numbers above the CO_2_ and CH_4_ emissions box plots are Spearman correlation coefficients between the respective gas emissions and the batch’s larval weight.

## Discussion

There are no field studies on direct CO_2_ and CH_4_ emissions from soil-inhabiting Scarabaeidae larvae yet to which we can compare our data^**4**^, but both the field- and laboratory-measured emission rates fall within the range of emission rates known from the few available laboratory studies on Scarabaeidae larvae and other soil Arthropoda groups in temperate regions^**5,13,14**^, or field and laboratory studies on termites in temperate, subtropical and tropical regions^**4,15,16**^. However, our study demonstrates how careful we have to be in interpreting GHG emission rates derived from laboratory studies. It has been known for termites that emission rates can decline over the course of a laboratory experiment^**17**^. In addition, we show that such a trend can also coincide with a considerable reduction of the emission rate variability between individual larvae and a disappearance of correlations between emission rates and environmental variables in comparison to field measurements.

Large variations in emission rates and the use of larval biomass for emission rate upscaling are features well known from termite studies^**10,16,18,19**^. Our biomass-based European CO_2_ and CH_4_ emission estimates for *Melolontha* spp. larvae are two orders of magnitude lower than the corresponding emission estimates for termites in temperate regions^**18**^. Termites are considered as a significant, but quite small global GHG source with the majority of these emissions stemming from subtropical and tropical regions^**4,10,20**^. Greenhouse gas emissions of soil Arthropoda groups other than termites are seen as too low to significantly affect regional budgets^**4,5**^, which our study seems to confirm at the first glance. However, in contrast to many termite studies, we did not attempt to use the emission rates of a few species to infer the emissions for this entire Arthropoda group. Our estimates are strictly for the genus *Melolontha* only, and thus, show only a fraction of the potential GHG emissions of European Scarabaeidae larvae. Worldwide, larvae of several Scarabaeidae species are regarded as economically important pest insects^**21,22**^. Regionally, these pest insects can reach biomass levels comparable to or considerably higher than those used for upscaling termite GHG emissions^**23**^, but we have no estimates of the total biomass of soil-dwelling Scarabaeidae larvae in Europe. Furthermore, for CH_4_ it is important to differentiate between gross and net soil fluxes. Purely biomass-based CH_4_ emission rates like ours represent gross soil CH_4_ fluxes, as they do not account for simultaneously occurring gross CH_4_ consumption in soils^**11,18,20**^. Recent studies suggests that CH_4_ emissions of soil-inhabiting Scarabaeidae larvae can stimulate soil CH_4_ consumption, and thus, potentially increase the overall net CH_4_ sink capacity of soils^**24,25**^. For regional and global CH_4_ budgets, it might therefore be more important to quantify the effect of soil faunal CH_4_ emissions on the net soil CH_4_ flux, instead of just quantifying total soil faunal CH_4_ emissions.

Regarding N_2_O, the emission rates measured in this study were of the same magnitude as those observed from earthworms^**26**^. Earthworms are the only faunal group for which a considerable amount of literature on soil N_2_O emissions is available^**27**^. In their presence, emissions can increase by more than 40 % due to the activation of nitrate- and nitrite-reducing bacteria during earthworm gut passage^**28**^. It is unclear if Scarabaeidae larvae are capable to affect soil N_2_O emissions in a similar manner as our data base it too inconsistent and no other data are available yet.

Overall, our data show that Scarabaeidae larvae should not be neglected as sources of CO_2_, CH_4_, and N_2_O in soil GHG flux research. However, to assess the impact of Scarabeidae larvae on regional and global GHG budgets and to better understand seasonal and interannual variations in GHG emissions, including the possibility of increased CH_4_ consumption in soils, it is mandatory to gather more field data on emission rates and species-dependent spatial larval biomass distributions and activities. These are exactly the same challenges, which are known from termite GHG emission research^**18**^, but we have to address these challenges if we want to explicitly include these soil faunal groups in future ecosystem and Earth system models.

## Acknowledgements

This project has received funding from the European Union’s Horizon 2020 research and innovation programme under the Marie Sklodowska-Curie grant agreement No 703107. We would like to thank Frauke Dormann (Geisenheim University) for the gas chromatographic analysis of the gas samples. We would also like to thank the following persons for access to the collection sites, help during the excavation of larvae, and ecological insights into these animals: Annemarie Peters (Hochschule Geisenheim University), Rainer Hurling and Sabine Weldner (NW-FVA – Northwest German Forest Research Institute), Wolfgang Dieminger and Hermann Stolz (local administration Blaubeuren-Weiler), Jana Reetz and Matthias Inthachot (LTZ – Center for Agricultural Technology Augustenberg), and Anne-Katrin Möller (district administration Alb-Donau-Kreis).

## Author contributions

C.M.G. designed the study, performed the measurements and the data analysis, and wrote the manuscript. C.K. initiated the topic and co-wrote the manuscript.

## Methods

### Sampling sites and species

This study was conducted in central and southern Germany – a temperate climate region with average annual air temperatures between 8 and 12 °C and average annual precipitation between 600 and 1000 mm (reference period 1961 – 1990)^**29**^. Target species were the Common cockchafer (*Melolontha melolontha*) and the Forest cockchafer (*M. hippocastani*) because they have a soil pest status in Europe and they represent two different distribution patterns of Scarabaeidae larvae in soils. Due to the pest status, they are one of the few European Scarabaeidae species for which regular monitoring programs exist, and thus, knowledge on larval ecology is relatively good^**12,23**^. *M. melolontha* and *M. hippocastani* live three and four years as root-feeding larvae in soils, respectively, progressing through three larval instars before pupating and evolving into adult beetles. The majority of individuals of a local population is of the same age, and population sizes tend to increase with every completed life cycle. *M. melolontha* inhabits open landscapes (e.g. pastures, vegetable crops, orchards, vineyards) and feeds in the rhizosphere mainly at 0 – 10 cm soil depth, while *M. hippocastani* inhabits forests and has a wider vertical soil profile distribution (0 – 40 cm soil depth) following tree root distribution^**12,30,31**^. During the 2017 vegetation period, larvae were excavated at six different sites covering a wide range of larval developmental stages and environmental conditions. Those sites were a meadow with *Beauveria* spp. infestation (Site 1), a Christmas tree plantation (tree height < 1m) (Site 2), a meadow without *Beauveria* spp. infestation (Site 3), and three mixed deciduous forest sites (Sites 4 – 6) (see supplementary information S2 for more details).

### Soil excavations

At each site, two to four randomly chosen plots with an area of 50 cm x 50 cm were carefully excavated by hand to a depth of ∼50 cm depending on site conditions. For each excavated larva, the following properties were recorded: excavation depth, weight, species and instar. The larvae were immediately subjected to gas emission measurements (see next paragraph). For species and instar identification, we relied on local expert opinion, but even for experts it was not always possible to distinguish between the two *Melolontha* species in the field. The adult beetles are easily identifiable as either *M. melolontha* or *M. hippocastani*; however, the larvae themselves can only be identified to the genus level based on morphological features alone^**32**^. For each plot, soil temperature and soil moisture were measured at the soil surface (0 – 5 cm depth) and at the plot’s final excavation depth (HH2 moisture meter with WET sensor, Delta-T Devices Ltd., Cambridge, United Kingdom). Air temperature and air pressure were measured with a handheld weather station (SM-28 Skymate PRO, WeatherHawk, Logan, UT, USA) at ∼2 m height at each plot.

### Gas emission measurements

Immediately following soil excavation, the larvae were individually placed in 110 ml glass tubes, which were sealed air-tight with butyl rubber stoppers. The larvae were incubated in the field for about an hour and a blank was included at each plot. No soil particles adhered to the larvae’s skin, but larvae could defecate during incubation. At the end of the incubation period, 25 ml of air were extracted from each glass tube with a plastic syringe and transferred to evacuated 12 ml glass vials sealed with grey chlorobutyl rubber septa (Labco Exetainers 839W, Labco Limited, Lampeter, United Kingdom), while control vacutainers were filled with ambient air during field incubations. At site 3, in addition to 18 larvae incubated directly in the field, 65 larvae were excavated and transferred to the laboratory, instead of being field-incubated. These larvae were kept individually in small plastic containers filled with ∼250 ml soil from the excavation site, and were supplied with ample amounts of fresh grass roots and carrot slices as food sources. Storage temperature was 18 °C and soils were sprayed with tab water once per week to keep them moist. After 7 days, 39 of these larvae were incubated in the laboratory (incubation temperature 24 °C) following the same protocol as in the field. After another 11 days, the remaining 26 larvae were incubated for gas sample collection. Gas samples were analysed with a SRI 8610C gas chromatograph with autosampler (SRI Instruments Europe GmbH, Bad Honnef, Germany) equipped with a flame ionisation detector (FID) coupled to a methanizer for CO_2_ and CH_4_ measurements, and an electron capture detector (ECD) for N_2_O measurements. Each detector was equipped with a Porapak Q pre-column (stainless steel tubing, length 1 m, 1/8 in. OD, 2 mm ID) and a Hayesep D column (stainless steel tubing, length 3 m, 1/8 in. OD, 2 mm ID). Column oven and detector temperatures were 70 °C and 330 °C, respectively. Nitrogen 5.0 (N_2_) was supplied as carrier gas at a pressure of 20 PSI. The ECD was additionally supplied with a make-up gas (4.5 % CO_2_ in N_2_ 5.0)^**33**^ at a pressure of 3 PSI. Due to temporary ECD failure, no N_2_O data are available for site 1 (see supplemental information S2). Peak integrations in the chromatograms were performed with PeakSimple Version 4.39 (6 channel) (SRI Instruments). Calibration curves were recorded with a 4-point standard gas series ranging from 304.0 to 3999.6 ppm CO_2_, 1.02 to 20.9 ppm CH_4_, and 248.4 to 15100 ppb N_2_O, respectively.

### Data processing and analysis

For each site, the CO_2_, CH_4_, and N_2_O emissions of each larva were quantified by subtracting the respective average incubation blank gas concentration value from each larval sample gas concentration value, applying the Ideal Gas Law^**34**^ and normalizing by incubation time. The relative error for each gas emission value was estimated via error propagation assuming the following random errors: 2 Kelvin for air temperature, 10 kPa for air pressure, and 0.01 L for the incubation volume. The random error for the blank-corrected larval sample gas concentration values (CO_2_ and CH_4_ in ppm, N_2_O in ppb) was the propagated error of the uncorrected larval sample gas concentration value and the incubation blank gas concentration value uncertainties derived from the gas chromatographic calibration curves. The relative propagated error for the larval CO_2_ and CH_4_ emission estimates ranged between 13 and 16 %, apart from a few exceptions. The relative propagated error for the final larval N_2_O emission estimates varied widely as the majority of N_2_O emissions was not significant. Emissions were classified as non-significant when the propagated error estimate exceeded the emission estimate.

A test for correlations between paired samples was performed on the entire pooled larval field dataset using Spearman’s *rho* (*r*_*s*_) statistic. Input variables were instar, larval excavation depth, larval weight (= biomass), larval abundance at the respective plot, individual CO_2_, CH_4_, and N_2_O emissions, air temperature, soil surface temperature and moisture, as well as soil temperature and moisture at the bottom of the respective excavated plot. For the comparison of larval field- and laboratory-measured emissions from Site 3, the same test statistic was also applied to the three flux data subsets (0, 7 and 18 days after excavation). The test was performed separately on each data subset with larval weight and larval CO_2_, CH_4_ and N_2_O emissions as input variables. Across-group comparisons on the data subsets were only carried out on larval weight to check for significant differences in mean biomass (Kruskal-Wallis rank sum test).

Larval CO_2_ and CH_4_ emissions were scaled up in two steps: from individual larvae to the plot level and from plot level to European level. Total larval emissions and larval biomass per plot were calculated by summing up the individual emissions or biomass and multiplying by four to scale to 1 m^2^. Larval biomass was subsequently used as independent variable in linear regression analysis for modelling m^2^-level larval CO_2_ and CH_4_ emissions. The obtained plot level larval CO_2_ and CH_4_ emissions were upscaled for 6 months per year (excluding larval winter rest) with the available literature data on European land area colonised by *Melolontha* spp. to derive a first rough annual CO_2_ and CH_4_ emission range estimate for Europe. A more precise upscaling to the European level is not yet possible, due to lack of field data. It needs to be considered that the available literature values on colonised land area are likely very conservative^**34**^. Due to their scattered occurrence, N_2_O emissions were not upscaled.

All test statistics and regression analysis were performed with the software R (version 3.4.3)^**35**^. In addition to the software’s standard library, the function ‘chart.Correlation’ (package: PerformanceAnalytics)^**36**^ was used.

