## Supplementary information S1 and S3 for "Scarabaeidae larvae are neglected greenhouse gas sources in soils"

### Supplementary information for manuscript *Scarabaeidae larvae are neglected greenhouse gas sources in soils*

Carolyn-Monika Görres

25 Juli 2019

#### S1 - Analysis of larval field emissions (on the level of the individual larvae)

A test for correlations between paired samples was performed on the entire pooled larval field dataset using Spearman's rho statistic. The following plot shows the correlations between $CO_2$ emission (mg $h^{-1}$ $larva^{-1}$), $CH_4$ emission (µg $h^{-1}$ $larva^{-1}$), larval biomass (g), and larval excavation depth (cm below soil surface).

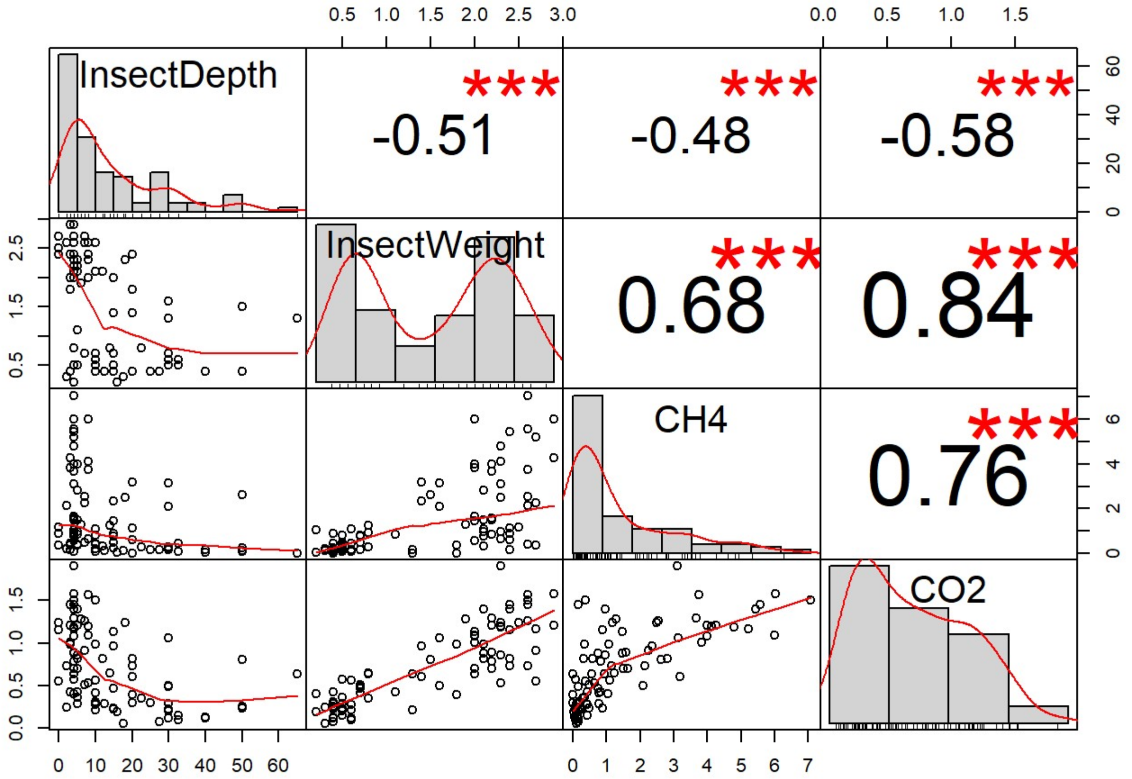


#### S3 - Regression analysis of cumulated larval $CO_2$ and $CH_4$ emissions estimates (plot level)

This section shows the results of the regression analysis which are shown in abbreviated form in Figure 2 in the manuscript. Cumulated larval $CO_2$ and $CH_4$ emission estimated from field measurements for each individual sampling plot were regressed on the total cumulated plot larval biomass (see Figure 2 in the manuscript).

This is the R result output for the $CO_2$ emission estimates.

```
##
## Call:
## lm(formula = Fig2Data$CO2plot ~ Fig2Data$PlotGrubBiomass)
##
## Residuals:
##     Min      1Q  Median      3Q     Max
## -6.8862 -1.4311  0.0607  1.7399  5.1025
##
## Coefficients:
##                           Estimate Std. Error t value Pr(>|t|)
## (Intercept)               -0.87828    1.45928  -0.602    0.558
## Fig2Data$PlotGrubBiomass  0.50868    0.03531  14.406 2.27e-09 ***
## ---
## Signif. codes:  0 '***' 0.001 '**' 0.01 '*' 0.05 '.' 0.1 ' ' 1
##
## Residual standard error: 3.462 on 13 degrees of freedom
## Multiple R-squared:  0.9411, Adjusted R-squared:  0.9365
## F-statistic: 207.5 on 1 and 13 DF,  p-value: 2.27e-09
```

This is the R result output for the $CH_4$ emission estimates.

```
##
## Call:
## lm(formula = Fig2Data$CH4plot ~ GrubBiomassSqr)
##
## Residuals:
##      Min       1Q   Median       3Q      Max
## -26.1256  -8.9636   0.1881   7.7654  24.7533
##
## Coefficients:
##                Estimate Std. Error t value Pr(>|t|)
## (Intercept)    0.578919   4.339499   0.133    0.896
## GrubBiomassSqr 0.019782   0.001538  12.864 9.04e-09 ***
## ---
## Signif. codes:  0 '***' 0.001 '**' 0.01 '*' 0.05 '.' 0.1 ' ' 1
##
## Residual standard error: 13.38 on 13 degrees of freedom
## Multiple R-squared:  0.9272, Adjusted R-squared:  0.9216
## F-statistic: 165.5 on 1 and 13 DF,  p-value: 9.037e-09
```
